# Expanding xylose metabolism in yeast for plant cell wall conversion to biofuels

**DOI:** 10.1101/007807

**Authors:** Xin Li, Vivian Yaci Yu, Yuping Lin, Kulika Chomvong, Raíssa Estrela, Julie M. Liang, Elizabeth A. Znameroski, Joanna Feehan, Soo Rin Kim, Yong-Su Jin, N. Louise Glass, Jamie H. D. Cate

**Author notes:** Correspondence should be addressed to J.H.D.C.

## Abstract

Sustainable biofuel production from renewable biomass will require the efficient and complete use of all abundant sugars in the plant cell wall. Using the cellulolytic fungus *Neurospora crassa* as a model, we identified a xylodextrin transport and consumption pathway required for its growth on hemicellulose. Successful reconstitution of this xylodextrin utilization pathway in *Saccharomyces cerevisiae* revealed that fungal xylose reductases act as xylodextrin reductases, and together with two hydrolases, generate intracellular xylose and xylitol. Xylodextrin consumption using xylodextrin reductases and tandem intracellular hydrolases greatly expands the capacity of yeasts to use plant cell wall-derived sugars, and should be adaptable to increase the efficiency of both first-generation and next-generation biofuel production.

---

The biological production of biofuels and renewable chemicals from plant biomass requires an economic way to convert complex carbohydrate polymers from the plant cell wall into simple sugars that can be fermented by microbes (Carroll and Somerville 2009, Chundawat, Beckham et al. 2011). In current industrial methods, cellulose and hemicellulose, the two major polysaccharides found in the plant cell wall (Somerville, Bauer et al. 2004), are generally processed into monomers of glucose and xylose, respectively (Chundawat, Beckham et al. 2011). However, in addition to harsh pretreatment of biomass, large quantities of cellulase and hemicellulase enzyme cocktails are required to release monosaccharides from plant cell walls, posing unsolved economic and logistical challenges (Lynd, Weimer et al. 2002, Himmel, Ding et al. 2007, Jarboe, Zhang et al. 2010, Chundawat, Beckham et al. 2011). The bioethanol industry currently uses the yeast *Saccharomyces cerevisiae* to ferment sugars derived from cornstarch or sugarcane into ethanol (Hong and Nielsen 2012), but *S. cerevisiae* requires substantial engineering to be able to ferment sugars derived from plant cell walls such as cellobiose and xylose (Kuyper, Hartog et al. 2005, Jeffries 2006, Ha, Galazka et al. 2011, Hong and Nielsen 2012, Young, Tong et al. 2014).

In contrast to *S. cerevisiae*, many cellulolytic fungi including *Neurospora crassa* (Tian, Beeson et al. 2009) naturally grow well on both the cellulose and hemicellulose components of the plant cell wall. By using transcription profiling data (Tian, Beeson et al. 2009) and by analyzing growth phenotypes of *N. crassa* knockout strains, we identified separate pathways used by *N. crassa* to consume cellodextrins (Galazka, Tian et al. 2010) and xylodextrins released by its secreted enzymes (Figure 1A and **Figure 1—figure supplement 1**) (Method 2014). A strain carrying a deletion of a previously-identified cellodextrin transporter (CDT-2, NCU08114) (Galazka, Tian et al. 2010) was unable to grow on xylan (**Figure 1—figure supplement 2**), and xylodextrins remained in the culture supernatant (**Figure 1—figure supplement 3**). As a direct test of transport function of CDT-2, *S. cerevisiae* strains expressing *cdt-2* were able to import xylobiose, xylotriose and xylotetraose (**Figure 1—figure supplement 4**). Notably, *N. crassa* expresses a putative intracellular β-xylosidase, GH43-2 (NCU01900), when grown on xylan (Sun, Tian et al. 2012). Purified GH43-2 displayed robust hydrolase activity towards xylodextrins with a degree of polymerization (DP) spanning from 2 to 8, and with a pH optimum near 7 (**Figure 1—figure supplement 5**). The results with CDT-2 and GH43-2 confirm those obtained independently in (Cai, Gu et al. 2014). As with CDT-1, orthologues of CDT-2 are widely distributed in the fungal kingdom (Galazka, Tian et al. 2010), suggesting that many fungi consume xylodextrins derived from plant cell walls. Furthermore, as with intracellular β-glucosidases (Galazka, Tian et al. 2010), intracellular β-xylosidases are also widespread in fungi (Sun, Tian et al. 2012) (**Figure 1—figure supplement 6**).

**Figure 1.**
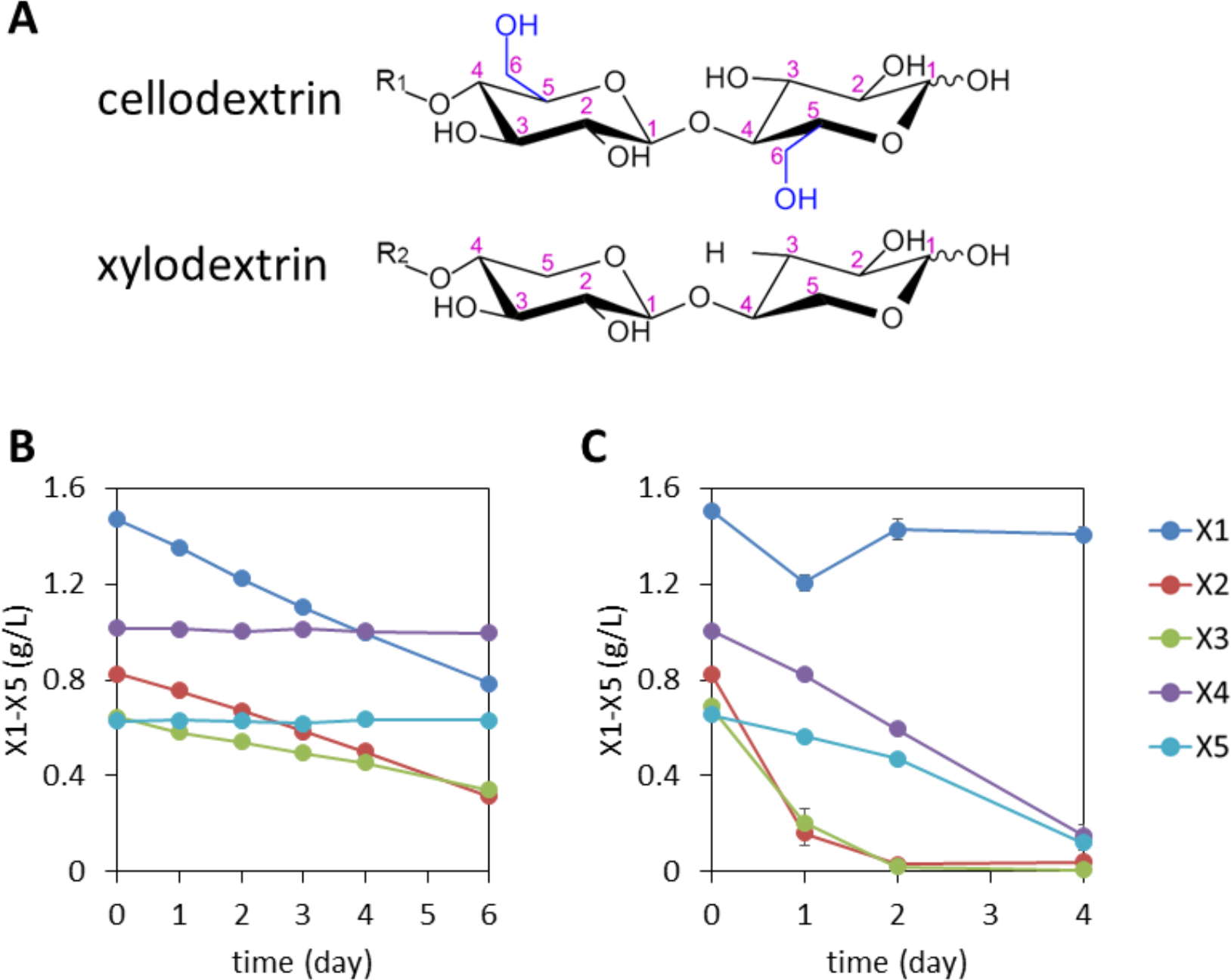
Consumption of xylodextrins by engineered *S. cerevisiae*. (**A**) Two oligosaccharide components derived from the plant cell wall. Cellodextrins, derived from cellulose, are a major source of glucose. Xylodextrins, derived from hemicellulose, are a major source of xylose. The 6-methoxy group (blue) distinguishes glucose derivatives from xylose. R_1_, R_2_ = H, cellobiose or xylobiose; R_1_ = β-1,4-linked glucose monomers in cellodextrins of larger degrees of polymerization; R_2_ = β-1,4-linked xylose monomers in xylodextrins of larger degrees of polymerization. (**B**) Xylose and xylodextrins remaining in a culture of *S. cerevisiae* grown on xylose and xylodextrins, and expressing an XR/XDH xylose consumption pathway, CDT-2, and GH43-2, with a starting cell density of OD600 = 1 under aerobic conditions. (**C**) Xylose and xylodextrins in a culture as in (**B**) but with a starting cell density of OD600 = 20. In both panels, the concentrations of xylose (X1) and xylodextrins with higher DPs (X2-X5) remaining in the culture broth after different periods of time are shown. All experiments were conducted in biological triplicate, with error bars representing standard deviations.

Cellodextrins and xylodextrins derived from plant cell walls are not catabolized by wild-type *S. cerevisiae* (Matsushika, Inoue et al. 2009, Galazka, Tian et al. 2010, Young, Lee et al. 2010). Reconstitution of a cellodextrin transport and consumption pathway from *N. crassa* in *S. cerevisiae* enabled this yeast to ferment cellobiose (Galazka, Tian et al. 2010). We therefore reasoned that expression of a functional xylodextrin transport and consumption system from *N. crassa* might further expand the capabilities of *S. cerevisiae* to utilize plant-derived xylodextrins as a source of carbon. Previously, *S. cerevisiae* was engineered to consume xylose by introducing xylose isomerase (XI), or by introducing xylose reductase (XR) and xylitol dehydrogenase (XDH) (Jeffries 2006, Matsushika, Inoue et al. 2009). To test whether *S. cerevisiae* could utilize xylodextrins, a *S. cerevisiae* strain was engineered with the XR/XDH pathway derived from *Scheffersomyces stipitis*–similar to the one utilized by *N. crassa* (Sun, Tian et al. 2012)–and a xylodextrin transport and consumption pathway from *N. crassa*. The xylose utilizing yeast expressing CDT-2 along with the intracellular β- xylosidase GH43-2 was able to directly utilize xylodextrins with DPs of 2 or 3 (Figure 1B and **Figure 1—figure supplement 7**).

Notably, although high cell density cultures of the engineered yeast were capable of consuming xylodextrins with DPs up to 5, xylose levels remained high (Figure 1C), suggesting the existence of severe bottlenecks in the engineered yeast. These results mirror those of a previous attempt to engineer *S. cerevisiae* for xylodextrin consumption, in which xylose was reported to accumulate in the culture medium (Fujii, Yu et al. 2011). Analyses of the supernatants from cultures with xylodextrins surprisingly revealed that the xylodextrins were converted into xylosyl-xylitol oligomers, a set of previously unknown compounds, by the engineered yeast, rather than hydrolyzed to xylose and consumed (Figure 2A **and Figure 2—figure supplement 1**). The resulting xylosyl-xylitol oligomers were effectively dead-end products that could not be metabolized further.

**Figure 2.**
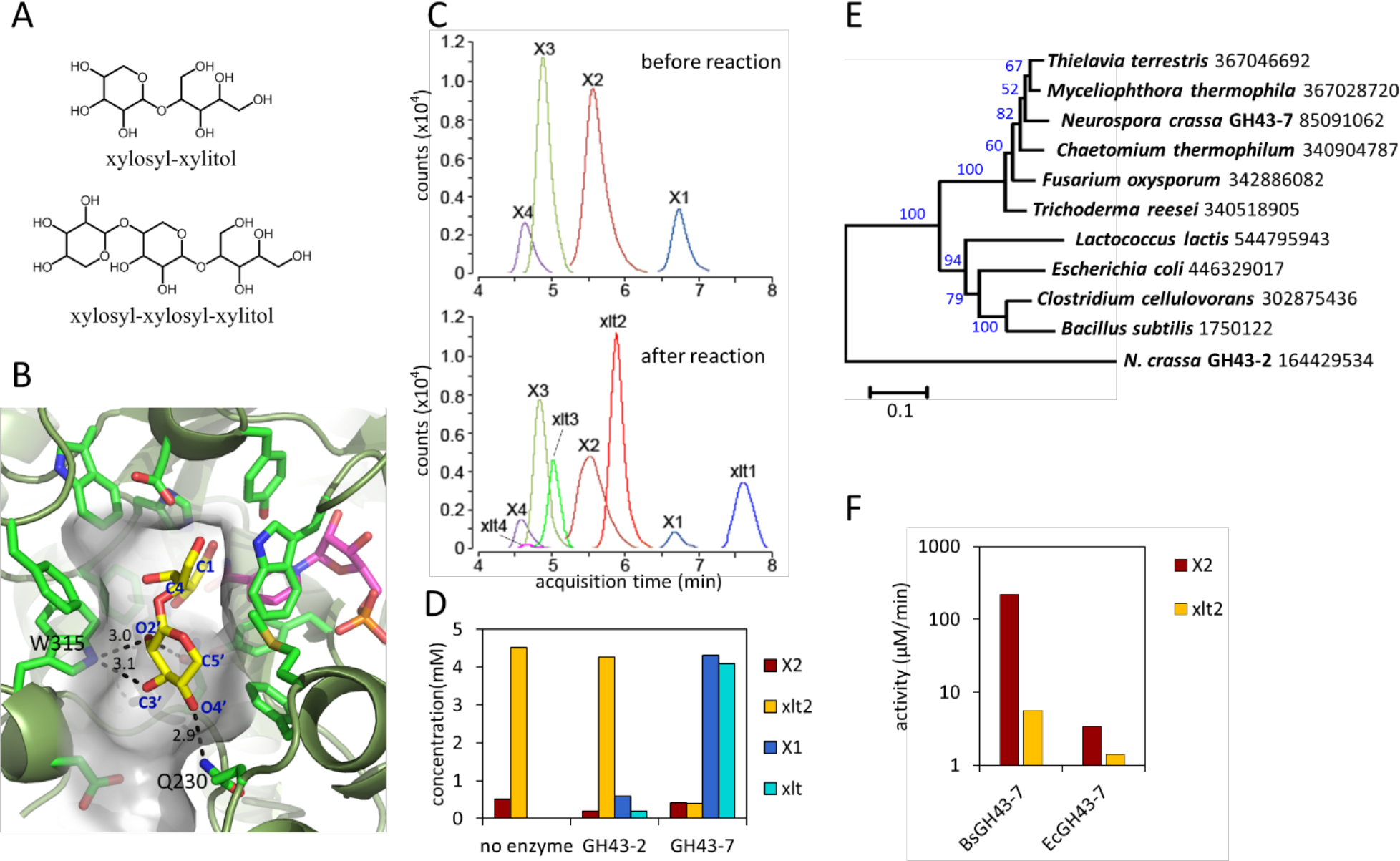
Production and enzymatic breakdown of xylosyl-xylitol. (**A**) Structures of xylosyl-xylitol and xylosyl-xylosyl-xylitol. (**B**) Computational docking model of xylobiose to *Ct*XR, with xylobiose in yellow, NADH cofactor in magenta, protein secondary structure in dark green, active site residues in bright green and showing side-chains. Part of the *Ct*XR surface is shown to depict the shape of the active site pocket. Black dotted lines show predicted hydrogen bonds between *Ct*XR and the non-reducing end residue of xylobiose. (**C**) Production of xylosyl-xylitol oligomers by *N. crassa* xylose reductase, XYR-1. Xylose, xylodextrins with DP of 2 to 4, and their reduced products are labeled X1-X4 and xlt1-xlt4, respectively. (**D**) Hydrolysis of xylosyl-xylitol by GH43-7. A mixture of 0.5 mM xylobiose and xylosyl-xylitol was used as substrates. Concentration of the products and the remaining substrates are shown after hydrolysis. (**E**) Phylogeny of GH43-7. *N. crassa* GH43-2 was used as an outgroup. 1000 bootstrap replicates were performed to calculate the supporting values shown on the branches. The scale bar indicates 0.1 substitutions per amino acid residue. The NCBI GI numbers of the sequences used to build the phylogenetic tree are indicated beside the species names. (**F**) Activity of two bacterial GH43-7 enzymes from *B. subtilis* (BsGH43-7) and *E. coli* (EcGH43-7).

Since the production of xylosyl-xylitol oligomers as intermediate metabolites has not been reported, the molecular components involved in their generation were examined. To test whether the xylosyl-xylitol oligomers resulted from side reactions of xylodextrins with endogenous *S. cerevisiae* enzymes, we used two separate yeast strains in a combined culture, one containing the xylodextrin hydrolysis pathway composed of CDT-2 and GH43-2, and the second engineered with the XR/XDH xylose consumption pathway. The strain expressing CDT-2 and GH43-2 would cleave xylodextrins to xylose, which could then be secreted via endogenous transporters (Hamacher, Becker et al. 2002) and serve as a carbon source for the strain expressing the xylose consumption pathway (XR and XDH). By contrast, the engineered yeast expressing XR and XDH would only be capable of consuming xylose (Figure 1B). When co-cultured, these strains consumed xylodextrins without producing the xylosyl-xylitol byproduct (**Figure 2—figure supplement 2**). These results indicate that endogenous yeast enzymes and GH43-2 transglycolysis activity are not responsible for generating the xylosyl-xylitol byproducts, i.e. that they must be generated by the XR from *S. stipitis* (*Ss*XR).

Fungal xylose reductases such as *Ss*XR have been widely used in industry for xylose fermentation. However, the structural details of substrate binding to the XR active site have not been established. To explore the molecular basis for XR reduction of oligomeric xylodextrins, the structure of *Candida tenuis* xylose reductase (*Ct*XR) (Kavanagh, Klimacek et al. 2002), a close homologue of *Ss*XR, was analyzed. *Ct*XR contains an open active site cavity where xylose could bind, located near the binding site for the NADH co-factor (Kavanagh, Klimacek et al. 2002, Kratzer, Leitgeb et al. 2006). Notably, the open shape of the active site can readily accommodate the binding of longer xylodextrin substrates (Figure 2B). Using computational docking algorithms (Trott and Olson 2010), xylobiose was found to fit well in the pocket. Furthermore, there are no obstructions in the protein that would prevent longer xylodextrin oligomers from binding (Figure 2B).

We reasoned that if the xylosyl-xylitol byproducts are generated by fungal XRs like that from *S. stipitis*, similar side products should be generated in *N. crassa*, thereby requiring an additional pathway for their consumption. Consistent with this hypothesis, xylose reductase XYR-1 (NCU08384) from *N. crassa* also generated xylosyl-xylitol products from xylodextrins (Figure 2C). However, when *N. crassa* was grown on xylan, no xylosyl-xylitol byproduct accumulated in the culture medium (**Figure 1—figure supplement 3**). Thus, *N. crassa* presumably expresses an additional enzymatic activity to consume xylosyl-xylitol oligomers. Consistent with this hypothesis, a second putative intracellular β-xylosidase upregulated when *N. crassa* was grown on xylan, GH43-7 (NCU09625) (Sun, Tian et al. 2012), had weak β-xylosidase activity but rapidly hydrolyzed xylosyl-xylitol into xylose and xylitol (Figure 2D and **Figure 2—figure supplement 3**). The newly-identified xylosyl-xylitol-specific β-xylosidase GH43-7 is widely distributed in fungi and bacteria (Figure 2E), suggesting that it is used by a variety of microbes in the consumption of xylodextrins. Indeed, GH43-7 enzymes from the bacteria *Bacillus subtilis* and *Escherichia coli* cleave both xylodextrin and xylosyl-xylitol (Figure 2F).

We next tested whether integration of the complete xylodextrin consumption pathway would overcome the poor xylodextrin utilization by *S. cerevisiae* (Figure 1) (Fujii, Yu et al. 2011). When combined with the original xylodextrin pathway (CDT-2 plus GH43-2), GH43-7 enabled *S. cerevisiae* to grow more rapidly on xylodextrin (Figure 3A) and precluded accumulation of xylosyl-xylitol intermediates (Figure 3B to D and **Figure 3—figure supplement 1**). The presence of xylose greatly improved anaerobic fermentation of xylodextrins (Figure 3E, **Figure 2—figure supplement 2 and Figure 2—figure supplement 3**), indicating that metabolic sensing in *S. cerevisiae* with the complete xylodextrin pathway may require additional tuning (Youk and van Oudenaarden 2009) for optimal xylodextrin fermentation. Taken together, these results reveal the XR/XDH pathway widely used in engineered *S. cerevisiae* naturally has broad substrate specificity for xylodextrins, and complete reconstitution of the naturally occurring xylodextrin pathway is necessary to enable *S. cerevisiae* to efficiently consume xylodextrins.

**Figure 3.**
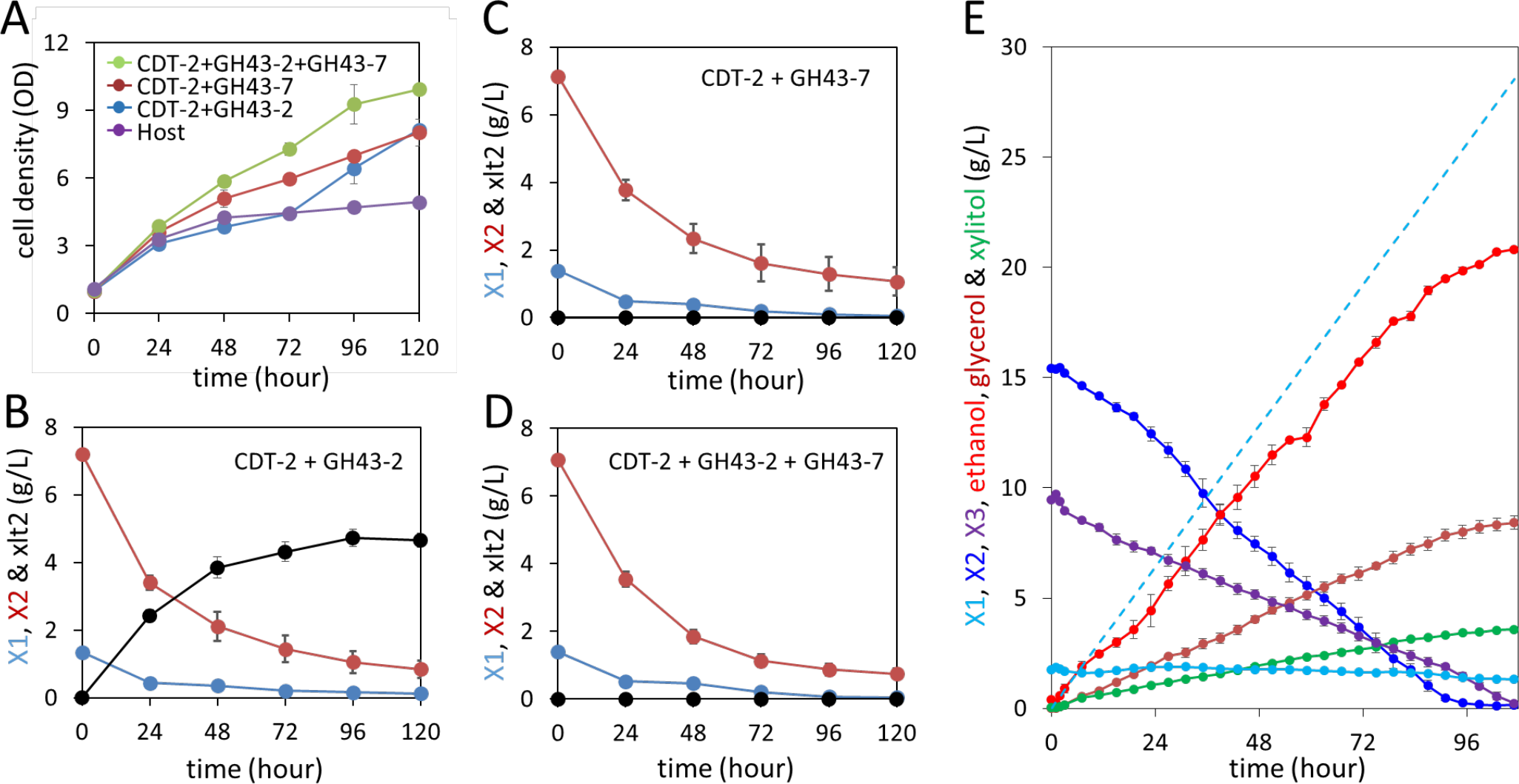
Aerobic consumption and anaerobic fermentation of xylodextrins with the complete xylodextrin pathway. (**A**)

Yeast growth curves with xylodextrin as the sole carbon source under aerobic conditions with a cell density at OD600 = 1. Yeast strain SR8U without plasmids, or transformed with plasmid expressing CDT-2 and GH43-2 (pXD8.4), CDT-2 and GH43-7 (pXD8.6) or all three genes (pXD8.7) are shown. (**B-D**) Xylobiose consumption with xylodextrin as the sole carbon source under aerobic conditions with a cell density of OD600 = 20. Xylosyl-xylitol accumulation was only observed in the SR8U strain bearing plasmid pXD8.4, i.e. lacking GH43-7. (**E**) Anaerobic fermentation of xylodextrins and xylose, in a fed-batch reactor. Strain SR8U expressing CDT-2, GH43-2, and GH43-7 (plasmid pXD8.7) was used at an initial OD600 of 20. Solid lines represent concentrations of compounds in the media. Blue dotted line shows the total amount of xylose added to the culture over time. Error bars represent standard deviations of biological triplicates (panels A-D) or duplicates (panel E).

Using yeast as a test platform, we identified a xylodextrin consumption pathway in *N. crassa* (Figure 4) that surprisingly involves a new metabolic intermediate likely to be widely produced in nature by many fungi and bacteria. In bacteria, xylosyl-xylitol may be generated by aldo-keto reductases known to possess broad substrate specificity (Barski, Tipparaju et al. 2008). The discovery of the xylodextrin consumption pathway along with cellodextrin consumption (Galazka, Tian et al. 2010) in cellulolytic fungi for the two major sugar components of the plant cell wall now provides many modes of engineering yeast to ferment plant biomass-derived sugars (Figure 4). Although XI could in principle replace the XR/XDH part of the pathway for producing intracellular xylulose, the XR/XDH pathway provides significant advantages in realistic fermentation conditions with sugars derived from hemicellulose. The breakdown of hemicellulose, which is acetylated (Sun, Tian et al. 2012), releases highly-toxic acetate, degrading the performance of *S. cerevisiae* fermentations (Bellissimi, van Dijken et al. 2009, Sun, Tian et al. 2012). The excess reducing power generated by the XR/XDH pathway, initially deemed a problem, can be exploited to drive acetate reduction, thereby detoxifying the fermentation medium and increasing ethanol production (Wei, Quarterman et al. 2013).

**Figure 4.**
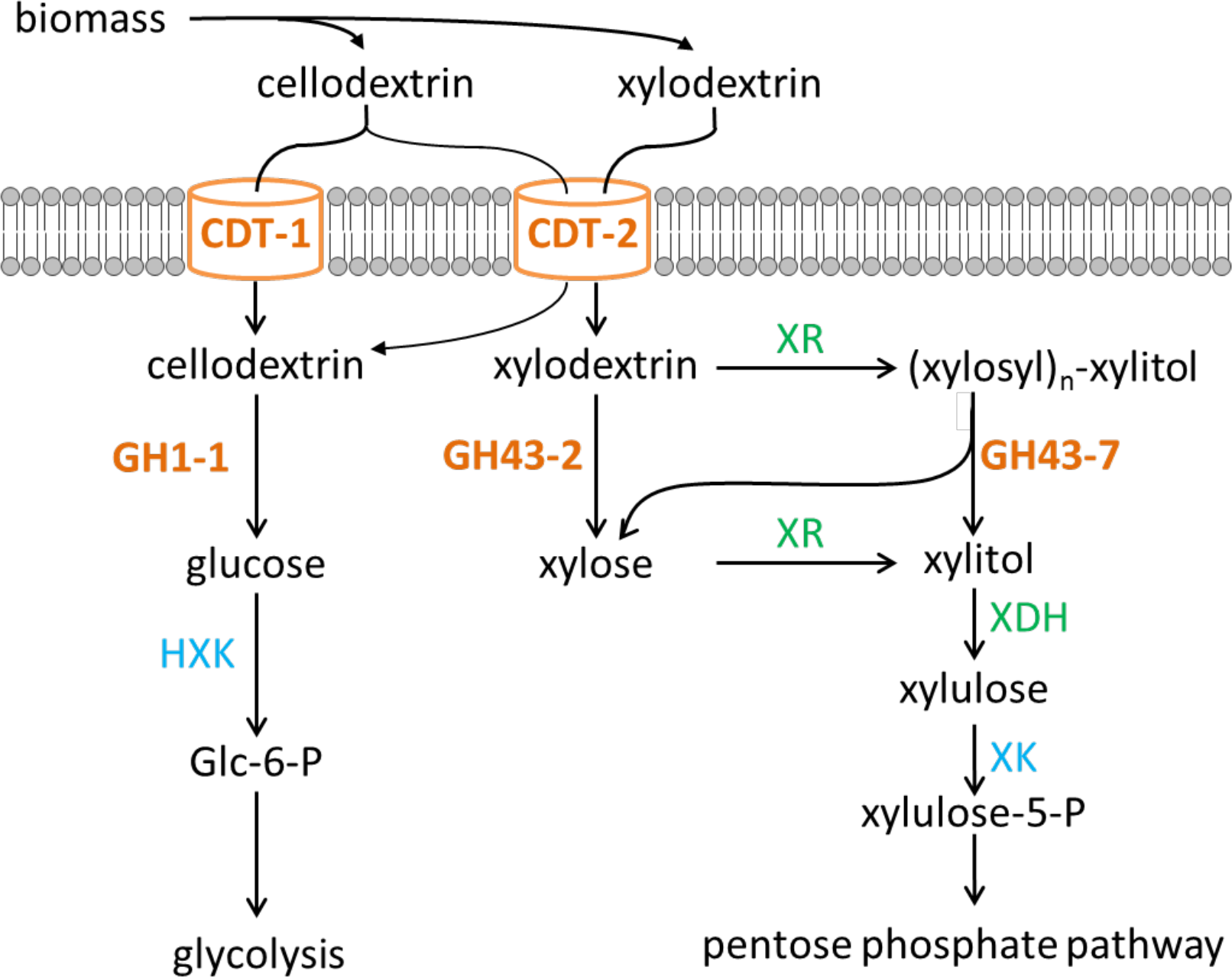
Two pathways of oligosaccharide consumption in *N. crassa* reconstituted in *S. cerevisiae*. Intracellular cellobiose utilization requires CDT-1 or CDT-2 along with β-glucosidase GH1-1 (Galazka, Tian et al. 2010), and enters glycolysis after phosphorylation by hexokinases (HXK) to form glucose-6-phosphate (Glc-6-P). Intracellular xylodextrin utilization also uses CDT-2, and requires the intracellular β-xylosidases GH43-2 and GH43-7. The resulting xylose can be assimilated through the pentose phosphate pathway consisting of xylose/xylodextrin reductase (XR), xylitol dehydrogenase (XDH), and xylulokinase (XK).

With optimization, the newly-identified xylodextrin consumption pathway provides new opportunities to expand first-generation bioethanol production from cornstarch or sugarcane to include hemicellulose from the plant cell wall. For example, xylodextrins and xylose released from the hemicellulose in sugarcane bagasse by using compressed hot water treatment (Hendriks and Zeeman 2009, Agbor, Cicek et al. 2011, Maria Evangelina Vallejos 2012) could be directly fermented by yeast engineered to consume xylodextrins. Xylodextrin consumption combined with cellodextrin consumption (Figure 4) should also improve next-generation biofuel production from lignocellulosic feedstocks under a number of pretreatment scenarios (Hendriks and Zeeman 2009, Maria Evangelina Vallejos 2012). These pathways could find widespread use to overcome remaining bottlenecks to fermentation of lignocellulosic feedstocks as a sustainable and economical source of biofuels and renewable chemicals.

## Materials and Methods

### Neurospora crassa strains

*N. crassa* strains obtained from the Fungal Genetics Stock Center (FGSC) (*22*) include the WT (FGSC 2489), and deletion strains for the two oligosaccharide transporters: NCU00801 (FGSC 16575) and NCU08114 (FGSC 17868).

### Neurospora crassa growth assay

Conidia were inoculated at a concentration equal to 10^6^ conidia per mL in 3 mL Vogel’s salts (Vogel 1956) with 2% wt/vol powdered *Miscanthus giganteus* (Energy Bioscience Institute, UC-Berkeley), Avicel PH 101 (Sigma), beechwood xylan (Sigma), or pectin (Sigma) in a 24-well deep-well plate. The plate was sealed with Corning™ breathable sealing tape and incubated at 25 °C in constant light and with shaking (200 rpm). Images were taken at 48 hours. Culture supernatants were diluted 200 times with 0.1 M NaOH before Dionex high-performance anion exchange chromatographic (HPAEC) analysis, as described below.

### Plasmids and yeast strains

Template gDNA from the *N. crassa* WT strain (FGSC 2489) and from the *S. cerevisiae* S288C strain was extracted as described in (http://www.fgsc.net/fgn35/lee35.pdf) and (*24*). Open reading frames (ORFs) of the β-xylosidase genes NCU01900 and NCU09652 (GH43-2 and GH43-7) were amplified from the *N. crassa* gDNA template. For biochemical assays, each ORF was fused with a C-terminal His_6_-tag and flanked with the *S. cerevisiae* P*_TEF1_* promoter and *CYC1* transcriptional terminator in the 2µ yeast plasmid pRS423 backbone. Plasmid pRS426_NCU08114 was described previously (Galazka, Tian et al. 2010). Plasmid pLNL78 containing the xylose utilization pathway (xylose reductase, xylitol dehydrogenase, and xylulose kinase) from *Scheffersomyces stipitis* was obtained from the lab of John Dueber (Latimer, Lee et al. 2014). Plasmid pXD2, a single-plasmid form of the xylodextrin pathway, was constructed by integrating NCU08114 and NCU01900 expression cassettes into pLNL78, using the In-Fusion cloning kit (Clontech). Plasmid pXD8.4 derived from plasmid pRS316 (Sikorski and Hieter 1989) was used to express CDT-2 and GH43-2, each from the P*_CCW12_* promoter. Plasmid pXD8.6 was derived from pXD8.4 by replacing the GH43-2 ORF with the ORF for GH43-7. pXD8.7 contained all three expression cassettes (CDT-2, GH43-2 and GH43-7) using the P*_CCW12_* promoter for each. *S. cerevisiae* strain D452-2 (*MAT***a** *leu2 his3 ura3 can1*) (Kurtzman 1994) and SR8U (the uracil autotrophic version of the evolved xylose fast utilization strain SR8) (Kim, Skerker et al. 2013) were used as recipient strains for the yeast experiments. The ORF for *N. crassa* xylose reductase (*xyr-1*, NcXR) was amplified from *N. crassa* gDNA and the introns were removed by overlapping PCR. XR ORF was fused to a C-terminal His_6_-tag and flanked with the *S. cerevisiae* P*_CCW12_* promoter and *CYC1* transcriptional terminator, and inserted into plasmid pRS313.

A list of the plasmids used in this study can be found in Table S1.

### Yeast cell-based xylodextrin uptake assay

*S. cerevisiae* was grown in an optimized minimum medium (oMM) lacking uracil into late log phase. The oMM contained 1.7 g/L YNB (Sigma, Y1251), 2-fold appropriate CSM dropout mixture, 10 g/L (NH_4_)_2_SO_4_, 1 g/L MgSO_4_.7H_2_O, 6 g/L KH_2_PO_4_, 100 mg/L adenine hemisulfate, 10 mg/L inositol, 100 mg/L glutamic acid, 20 mg/L lysine, 375 mg/L serine and 100 mM 4-morpholineethanesulfonic acid (MES), pH 6.0 (Lin, Chomvong et al. Submitted). Cells were then harvested and washed three times with assay buffer (5mM MES, 100 mM NaCl, pH 6.0) and resuspended to a final OD600 of 40. Substrate stocks were prepared in the same assay buffer at a concentration of 200 µM. Transport assays were initiated by mixing equal volumes of the cell suspension and the substrate stock. Reactions were incubated at 30 °C with continuous shaking for 30 minutes. Samples were centrifuged at 14,000 rpm at 4 °C for 5 minutes to remove yeast cells. 400 µL of each sample supernatant was transferred to an HPLC vial containing 100 µL 0.5 M NaOH, and the concentration of the remaining substrate was measured by HPAEC as described below.

### Enzyme purification

*S. cerevisiae* strains transformed with pRS423_GH43-2, pRS423_GH43-7, or pRS313_NcXR were grown in oMM lacking histidine with 2% glucose until late log phase before harvesting by centrifugation. *E. coli* strains BL21DE3 transformed with pET302_BsGH43-7 or pET302_EcGH43-7 were grown in TB medium, induced with 0.2 mM IPTG at OD600 of 0.8, and harvested by centrifugation 12 hours after induction. Yeast or *E. coli* cell pellets were resuspended in a buffer containing 50 mM Tris-HCl, 100 mM NaCl, 0.5 mM DTT, pH 7.4 and protease inhibitor cocktail (Pierce). Cells were lysed with an Avestin homogenizer, and the clarified supernatant was loaded onto a HisTrap column (GE Healthcare). His-tagged enzymes were purified with an imidazole gradient, buffer-exchanged into 20 mM Tris-HCl, 100 mM NaCl, pH 7.4, and concentrated to 5 mg/mL.

### Enzyme assays

For the β-xylosidase assay of GH43-2 with xylodextrins, 0.5 µM of purified enzyme was incubated with 0.1% in-house prepared xylodextrin or 1 mM xylobiose (Megazyme) in 1x PBS at 30 °C. Reactions were sampled at 30 min and quenched by adding 5 volumes of 0.1 M NaOH. The products were analyzed by HPAEC as described below. For pH profiling, acetate buffer at pH 4.0, 4.5, 5.0, 5.5, 6.0, and phosphate buffer at 6.5, 7.0, 7.5, 8 were added at a concentration of 0.1 M. For the β-xylosidase assay of GH43-2 and GH43-7 with xylosyl-xylitol, 10 µM of purified enzyme was incubated with 4.5 mM xylosyl-xylitol and 0.5 mM xylobiose in 20 mM MES buffer, pH = 7.0, and 1 mM CaCl_2_ at 30 °C. Reactions were sampled at 3 hours and quenched by heating at 99 °C for 10 min. The products were analyzed by ion-exclusion HPLC as described below.

For the xylose reductase assays of NcXR, 1 µM of purified enzyme was incubated with 0.06% xylodextrin and 2 mM NADPH in 1x PBS at 30 °C. Reactions were sampled at 30 min and quenched by heating at 99 °C for 10 min. The products were analyzed by LC-QToF as described below.

### Oligosacchride preparation

Xylodextrin was purchased from Cascade Analytical Reagents and Biochemicals or prepared according to published procedures (Akpinar, Erdogan et al. 2009) with slight modifications. In brief, 20 g beechwood xylan (Sigma-Aldrich) was fully suspended in 1000 mL water, to which 13.6 mL 18.4 M H_2_SO_4_ was added. The mixture was incubated in a 150 °C oil bath with continuous stirring. After 30 min, the reaction was poured into a 2L plastic container on ice, with stirring to allow it to cool. Then 0.25 mol CaCO_3_ was slowly added to neutralize the pH and precipitate sulfate. The supernatant was filtered and concentrated on a rotary evaporator at 50 °C to dryness. The in-house prepared xylodextrin contained about 30% xylose monomers and 70% oligomers. To obtain a larger fraction of short chain xylodextrin, the commercial xylodextrin was dissolved to 20% w/v and incubated with 2 mg/mL xylanase at 37 °C for 48 hours. Heat deactivation and filtration were performed before use.

Xylosyl-xylitol was purified from the culture broth of strain SR8 containing plasmids pXD8.4 in xylodextrin medium. 50 mL of culture supernatant was concentrated on a rotary evaporator at 50 °C to about 5 mL. The filtered sample was loaded on an XK 16/70 column (GE Healthcare) packed with Supelclean^TM^ ENVI-Carb^TM^ (Sigma-Aldrich) mounted on an ÄKTA Purifier (GE Healthcare). The column was eluted with a gradient of acetonitrile at a flow rate of 3.0 mL/min at room temperature. Purified fractions, verified by LC-MS, were pooled and concentrated. The final product, containing 90% of xylosyl-xylitol and 10% xylobiose, was used as the substrate for enzyme assays and as an HPLC calibration standard.

### Yeast cultures with xylodextrins

Yeast strains were pre-grown aerobically overnight in oMM medium containing 2% glucose, washed 3 times with water, and resuspended in oMM medium. For aerobic growth, strains were inoculated at a starting OD600 of 1.0 or 20 in 50 mL oMM medium with 3% w/v xylodextrins and cultivated in 250 mL Erlenmeyer flasks covered with 4 layers of miracle cloth, shaking at 220 rpm. At the indicated time points, 0.8 mL samples were removed and pelleted. 20 µL supernatants were analyzed by ion-exclusion HPLC to determine xylose, xylitol, glycerol, and ethanol concentrations. 25 µL of 1:200 diluted or 2 µL of 1:100 diluted supernatant was analyzed by HPAEC or LC-QToF, respectively, to determine xylodextrin concentrations.

### Anaerobic fermentation

Anaerobic fermentation experiments were performed in a 1 L stirred tank bioreactor (DASGIP Bioreactor system, Type DGCS4, Eppendorf AG), containing oMM medium with 3% w/v xylodextrins inoculated with an initial cell concentration of OD600 = 20. The runs were performed at 30 °C for 107 h. The culture was agitated at 200 rpm and purged constantly with 6 L/h of nitrogen. Xylose was fed continuously at 0.8 mL/h from a 25% stock. During the fermetnation, 3 mL cell-free samples were taken each 4 h with an autosampler through a ceramic sampling probe (Seg-Flow Sampling System, Flownamics). 20 µL of the supernatant fraction were analyzed by ion-exclusion HPLC to determine xylose, xylitol, glycerol, acetate and ethanol concentrations. 2 µL of 1:100 diluted supernatant was analyzed by LC-QToF to determine xylodextrin concentrations.

### Ion-exclusion HPLC analysis

Ion-exclusion HPLC was performed on a Shimadzu Prominence HPLC equipped with a refractive index detector. Samples were resolved on an ion exclusion column (Aminex HPX-87H Column, 300 x 7.8 mm, Bio-Rad) using a mobile phase of 0.01 N H_2_SO_4_ at a flow rate of 0.6 mL/min at 50 °C.

### HPAEC analysis

HPAEC analysis was performed on a ICS-3000 HPLC (Thermo Fisher) using a CarboPac PA200 analytical column (150 × 3 mm) and a CarboPac PA200 guard column (3 × 30 mm) at 30 °C. Following injection of 25 µL of diluted samples, elution was performed at 0.4 mL/min using 0.1 M NaOH in the mobile phase with sodium acetate gradients. For xylodextrin and xylosyl-xylitol separation, the acetate gradients were 0 mM for 1 min, increasing to 80 mM in 8 min, increasing to 300 mM in 1 min, keeping at 30 mM for 2 min, followed by re-equilibration at 0 mM for 3 min. Carbohydrates were detected using pulsed amperometric detection (PAD) and peaks were analyzed and quantified using the Chromeleon software package.

### Mass spectrometric analyses

All mass spectrometric analyses were performed on an Agilent 6520 Accurate-Mass Q-TOF coupled with an Agilent 1200 LC. Samples were resolved on a 100x 7.8 mm Rezex RFQ-Fast Fruit H+ 8% column (Phenomenex) using a mobile phase of 0.5% formic acid at a flow rate of 0.3 mL/min at 55 °C.

To determine the accurate masses of the unknown metabolites, 2 µL of 1:100 diluted yeast culture supernatant was analyzed by LC-QTOF. Nitrogen was used as the instrument gas. The source voltage (Vcap) was 3,000 V in negative ion mode, and the fragmentor was set to 100 V. The drying gas temperature was 300 °C; drying gas flow was 7 L/min; and nebulizer pressure was 45 psi. The ESI source used a separate nebulizer for the continuous, low-level introduction of reference mass compounds (112.985587, 1033.988109) to maintain mass axis calibration. Data was collected at an acquisition rate of 1 Hz from m/z 50 to 1100, and stored in centroid mode.

LC-MS/MS was performed to confirm the identity of xylosyl-xylitol and xylosyl-xylosyl-xylitol. The compound with a retention time (RT) of 5.8 min and m/z ratio of 283.103 and the compound with an RT of 4.7 min and m/z ratio of 415.15 were fragmented with collision energies of 10, 20 and 40 eV. MS/MS spectra were acquired, and the product ions were compared and matched to the calculated fragment ions generated by the Fragmentation Tools in ChemBioDraw Ultra v13.

To quantify the carbohydrates and carbohydrate derivatives in the culture, culture supernatants were diluted 100-fold in water and 2 µL was analyzed by LC-QToF. Spectra were imported to Qualtitative Analysis module of Agilent MassHunter Workstation software using m/z and retention time values obtained from the calibration samples to search for the targeted ions in the data. These searches generated extracted ion chromatograms (EICs) based on the list of target compounds. Peaks were integrated and compared to the calibration curves to calculate the concentration. Calibration curves were calculated from the calibration samples, prepared in the same oMM medium as all the samples, and curve fitting for each compound resulted in fits with R^2^ values of 0.999. 4-morpholineethanesulfonic acid (MES), the buffer compound in the oMM medium with constant concentration and not utilized by yeast, was used as an internal standard (IS) for concentration normalization.

## Acknowledgement

We thank L. Acosta-Sampson and A. Gokhale for helpful discussions; J. Dueber for xylose utilization pathway plasmids, Z. Baer, J. Kuchenreuthe and M. Maurer for helps in anaerobic fermentation, and S. Bauer and A. Ibañez Zamora for help with analytical methods. This work was supported by funding from the Energy Biosciences Institute (J.H.D.C., N.L.G and Y.S.J.), and by a predoctoral fellowship from CNPq and CAPES through the program “Ciência sem Fronteiras” (R. E.).

## Supplementary file

**Figure 1—figure supplement 1.**
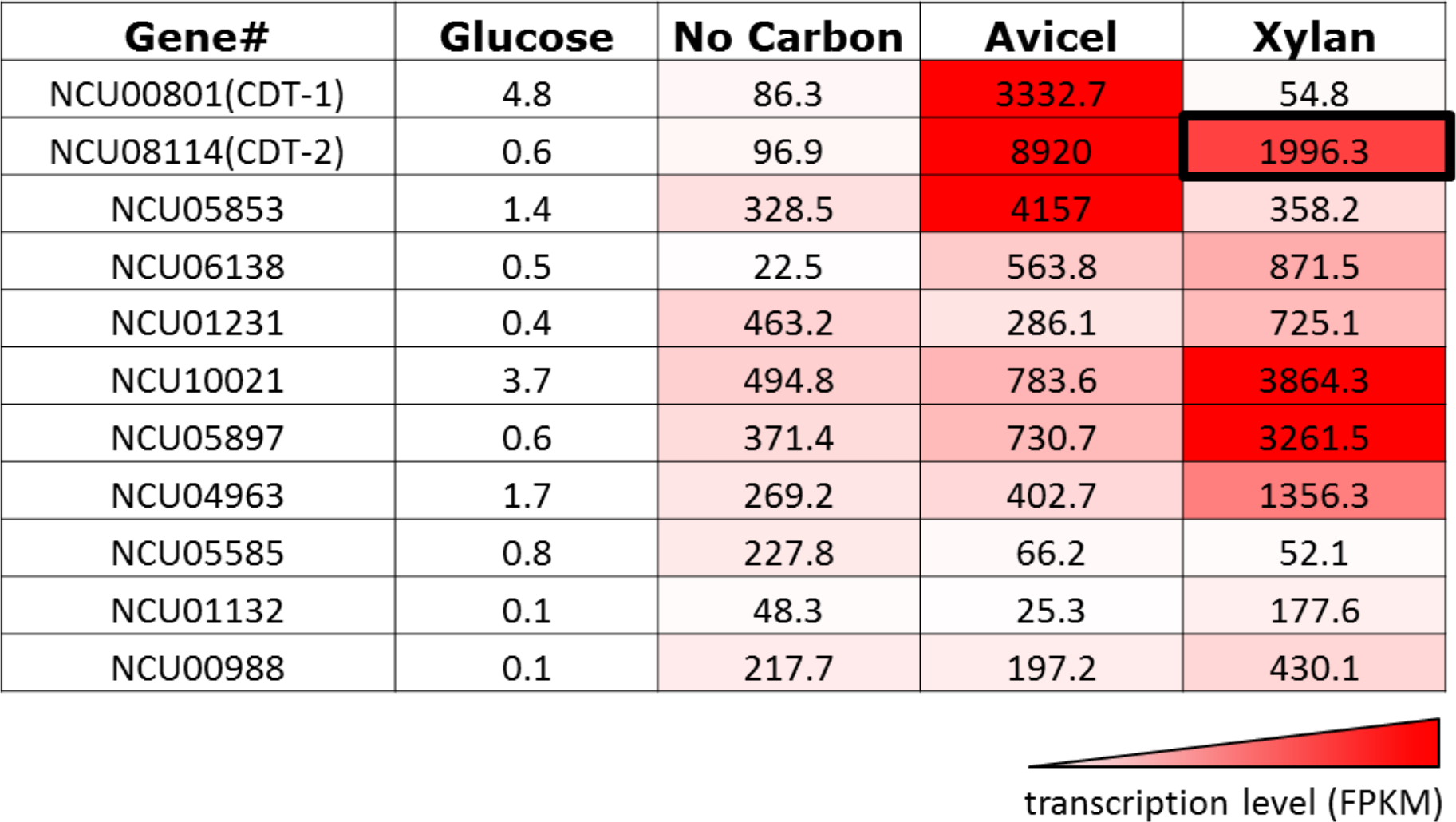
Transcriptional levels of transporters expressed in *N. crassa* grown on different carbon sources.

Transcript levels reported in fragments per kilobase per million reads (FPKM) are derived from experiments published in (Coradetti, Craig et al. 2012, Sun, Tian et al. 2012).

**Figure 1—figure supplement 2.**
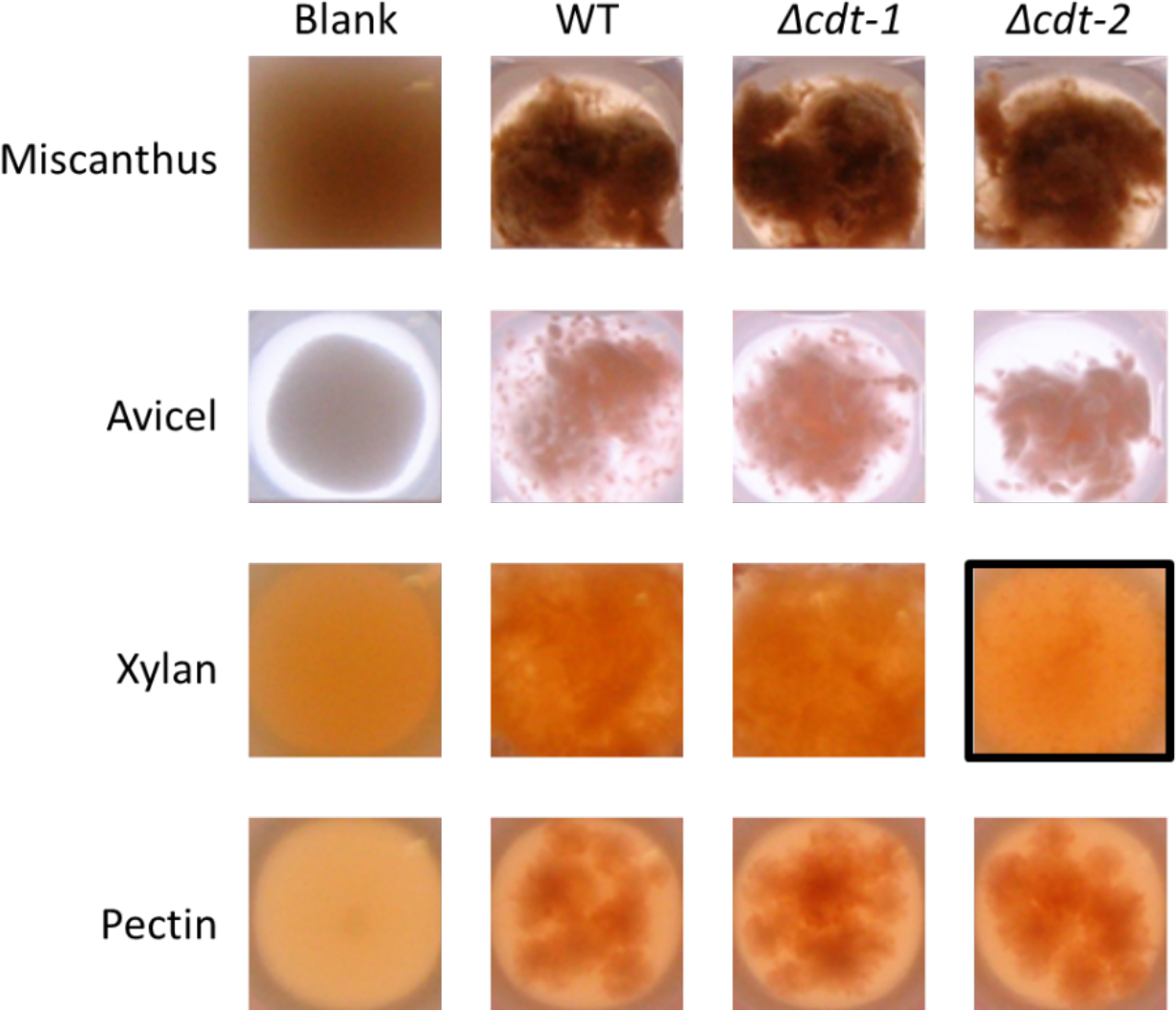
Growth of *N. crassa* strains on different carbon sources.

Wild type (WT) *N. crassa*, or *N. crassa* with deletions of transporters *cdt-1* (Δ*cdt-1*) or *cdt-2* (Δ*cdt-2*), were grown on *Miscanthus giganteus* plant cell walls, or purified plant cell wall components. Avicel is a form of cellulose derived from plant cell walls. The black box shows the severe growth phenotype of the Δ*cdt-2* strain grown on xylan medium.

**Figure 1—figure supplement 3.**
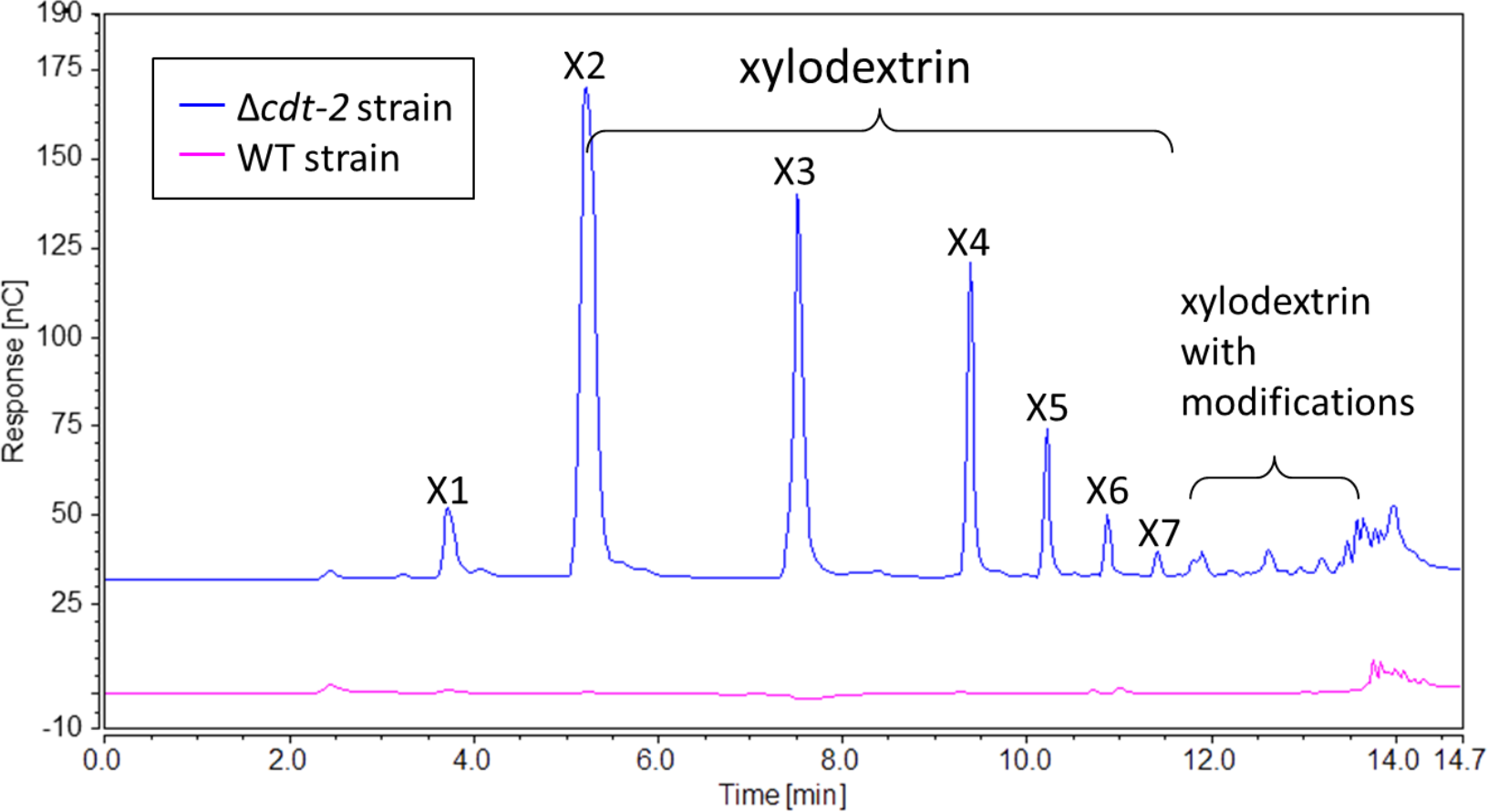
Xylodextrins in the xylan culture supernatant of the *N. crassa* Δ*cdt-2* strain.

25 µL of 1:200 diluted *N. crassa* xylan culture supernantant was analyzed by HPAEC on a CarboPac PA200 column. While no detectable soluble sugars were found in the culture supernatant of the wild-type strain (magenta line), the Δ*cdt-2* strain (blue line) left a high concentration of unmodified and modified xylodextrins in the culture supernatant. Little xylose was found, indicating xylose was transported by means of different transporters.

**Figure 1—figure supplement 4.**
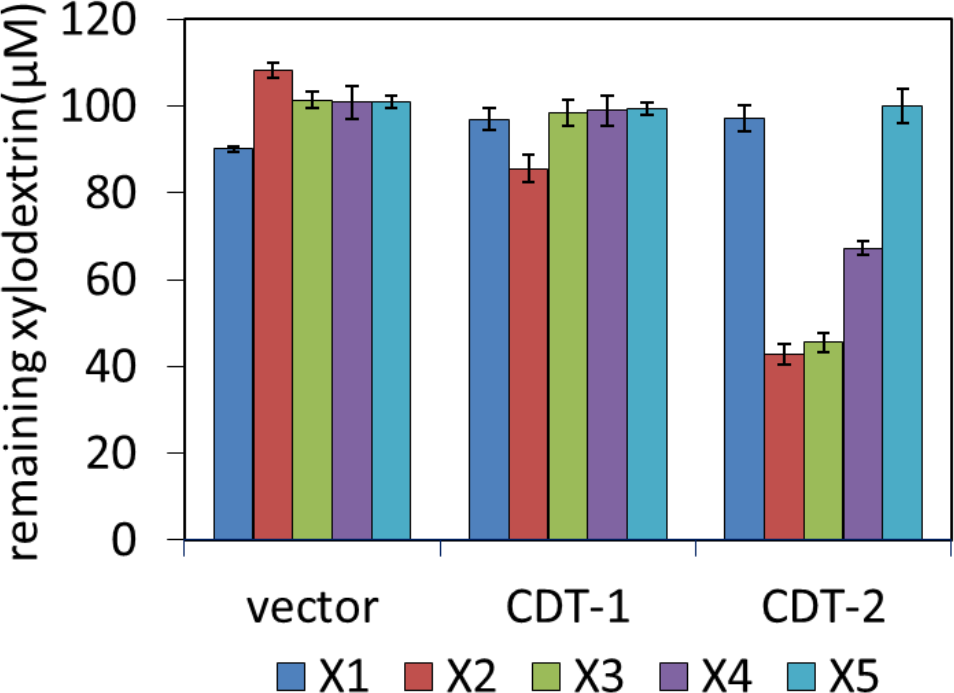
Transport of xylodextrins into the cytoplasm of *S. cerevisiae* strains expressing *N. crassa* transporters.

The starting xylodextrin concentration for each purified component was 100 µM. The remaining xylose (X1) and xylodextrins in the culture media are shown for experiments with *S. cerevisiae* harboring an empty expression plasmid (vector), or with *S. cerevisiae* individually expressing transporters CDT-1 or CDT-2. Xylodextrins used include xylobiose (X2), xylotriose (X3), xylotetraose (X4), and xylopentaose (X5). Error bars indicate standard deviations of biological triplicates.

**Figure 1—figure supplement 5.**
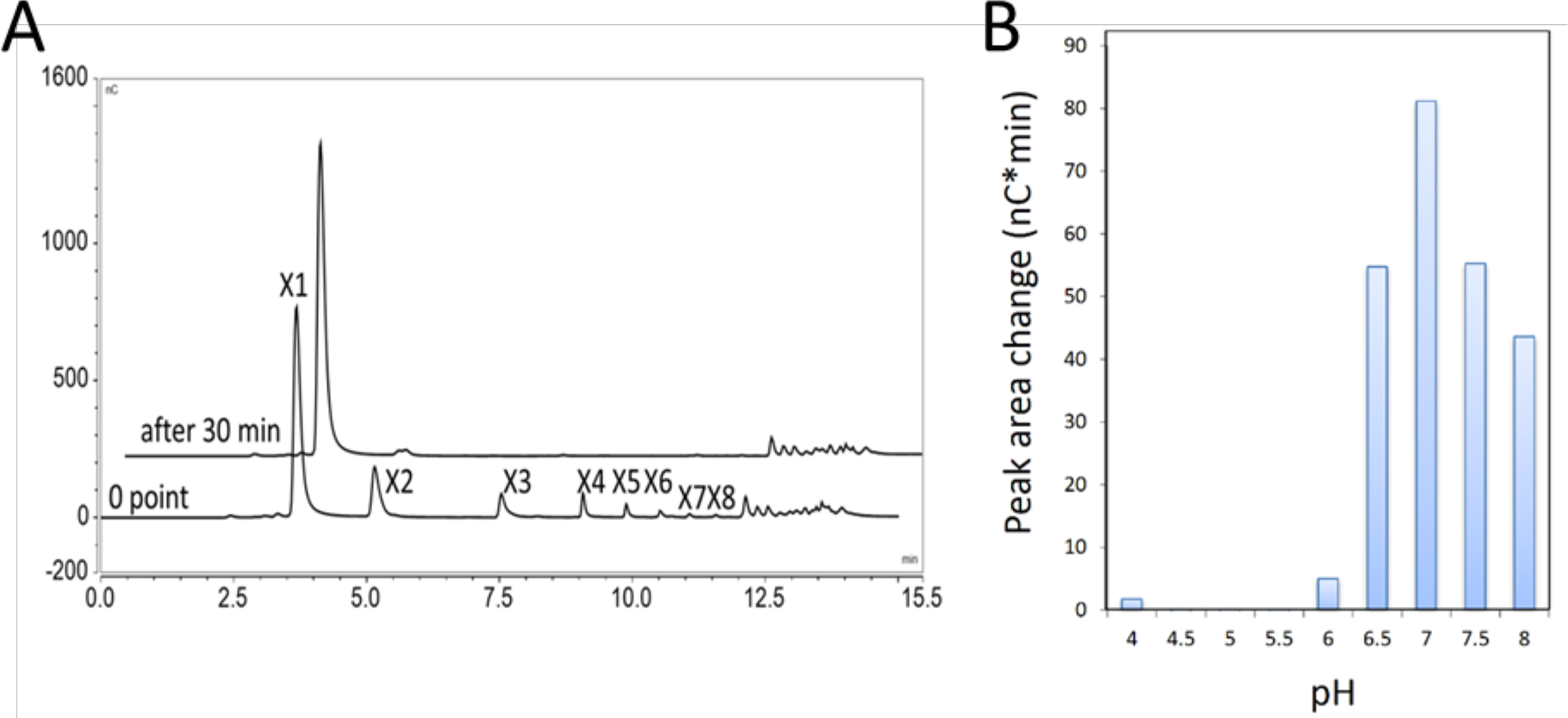
Xylobiase activity of the predicted β-xylosidase GH43-2.

(**A**) GH43-2 hydrolysis of xylodextrins with degrees of polymerization from at least 2-8. The 30 min chromatogram is offset for clarity. (**B**) The pH optimum of GH43-2, determined by measuring the extent of hydrolysis of xylobiose to xylose. The HPAEC chromatogram peak area change for xylose is shown.

**Figure 1—figure supplement 6.**
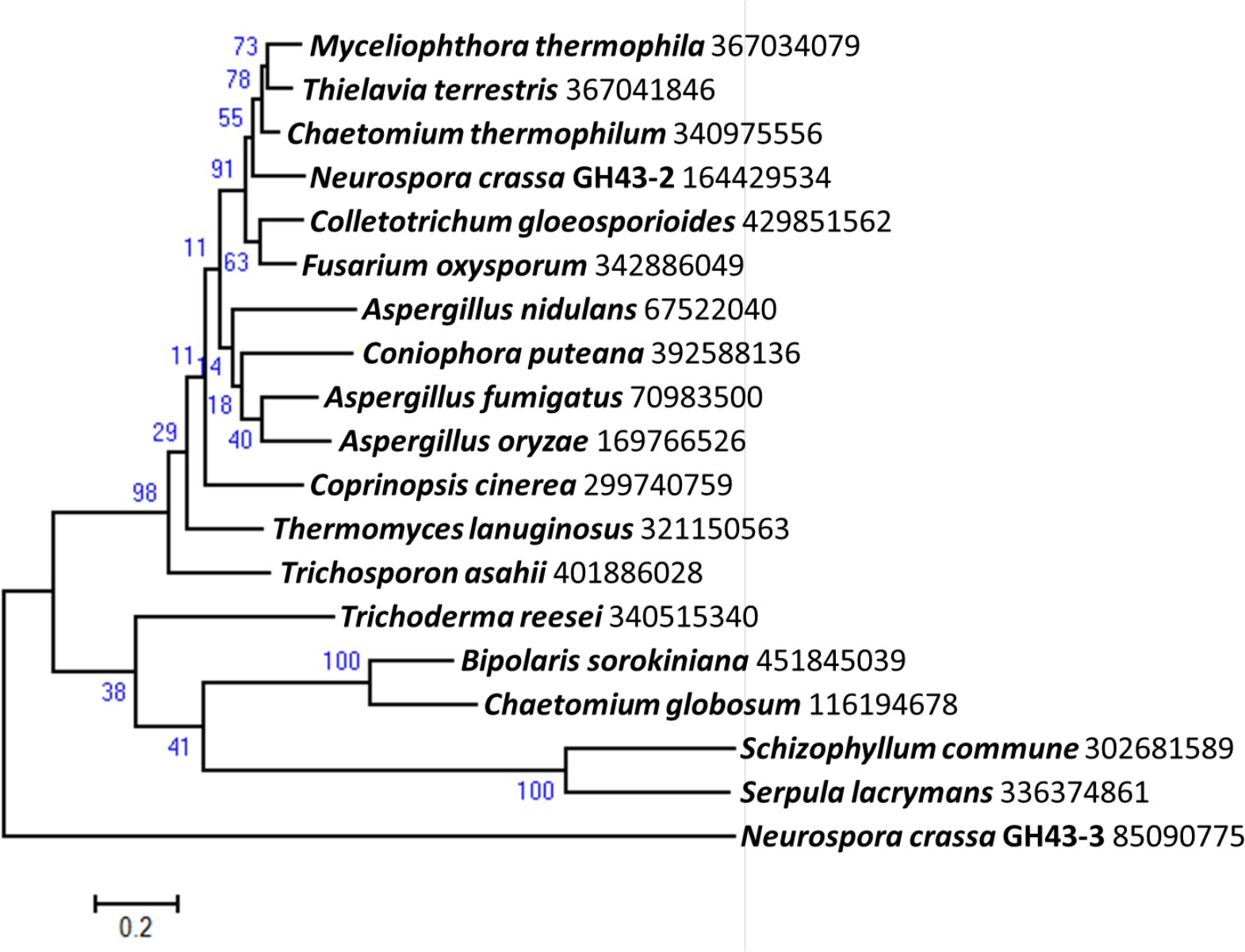
Phylogenetic distribution of predicted intracellular β-xylosidases GH43-2 in filamentous fungi.

Homologs of GH43-2 (NCU01900) were found with BLAST (Altschul, Madden et al. 1997) queries of respective sequence against NCBI protein database. Representative sequences from a diversified taxonomy were chosen and aligned with the MUSCLE algorithm (Edgar 2004). A maximum likelihood phylogenetic tree was calculated based on the alignment with the Jones-Taylor-Thornton model by using software MEGA v6.05 (Tamura, Stecher et al. 2013). Xylan-induced extracellular GH43-3 (NCU05965) was used as an outgroup. The NCBI GI numbers of the sequences used to build the phylogenetic tree were indicated besides the species names. 1000 bootstrap replicates were performed to calculate the supporting values shown on the branches. The scale bar indicates 0.2 substitutions per amino acid residue.

**Figure 1—figure supplement 7.**
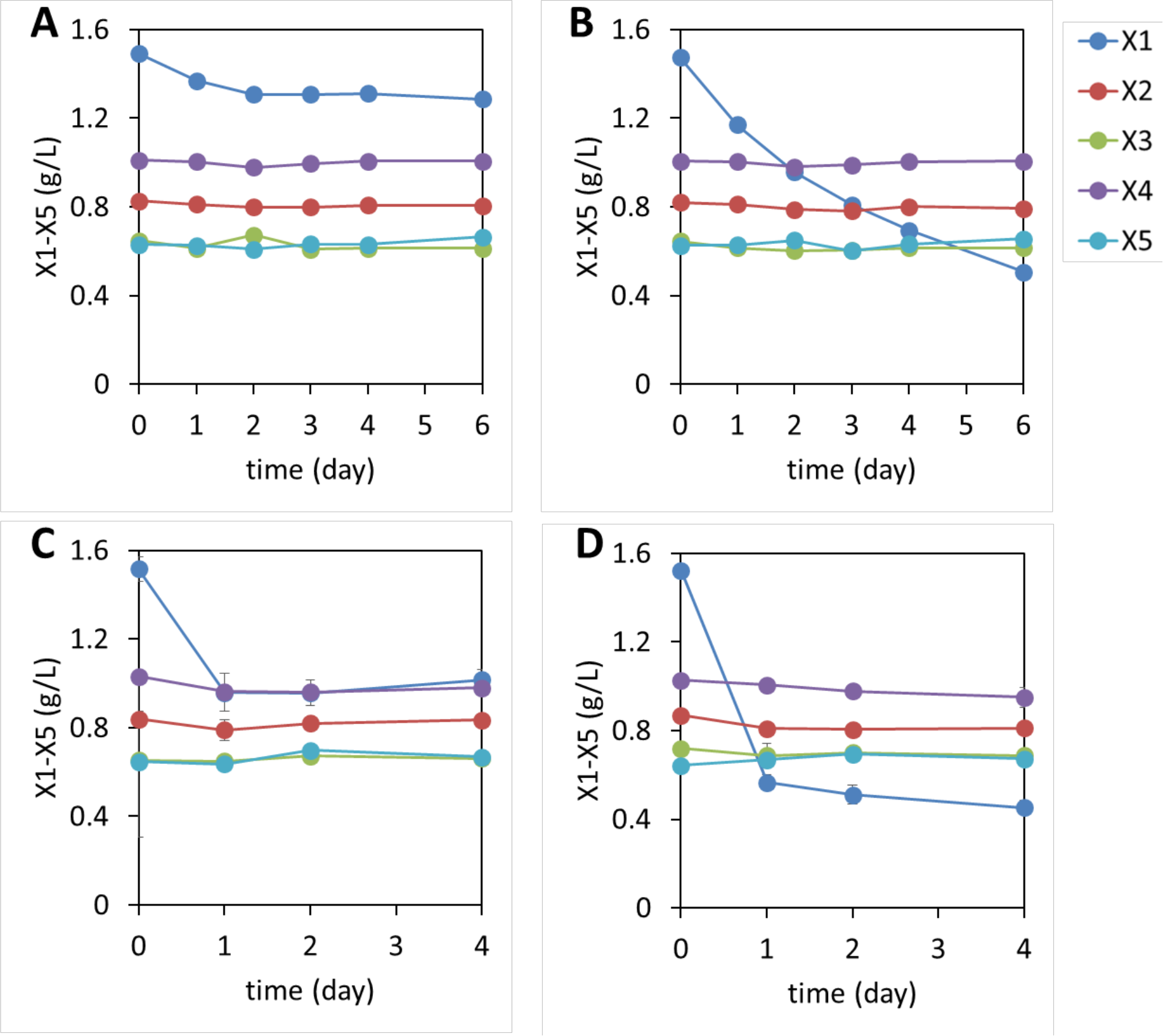
Xylodextrin consumption profiles of *S. cerevisiae* strains lacking the xylodextrin pathway.

Shown are the concentrations of the remaining sugars in the culture broth after different periods of time of (**A**) the WT D452-2 strain with starting cell density at OD600 = 1, (**B**) D452-2 with a *S. stipitis* xylose utilization pathway (plasmid pLNL78) with a starting cell density at OD600 = 1, (**C**) WT D452-2 strain with a starting cell density at OD600 = 20, and (**D**) D452-2 with a *S. stipitis* xylose utilization pathway (plasmid pLNL78) with a starting cell density at OD600 = 20. In all panels, xylose (X1), and xylodextrins of higher DPs (X2-X5) are shown. Error bars represent standard deviations of biological triplicates (panels A-D).

**Figure 2—figure supplement 1.**
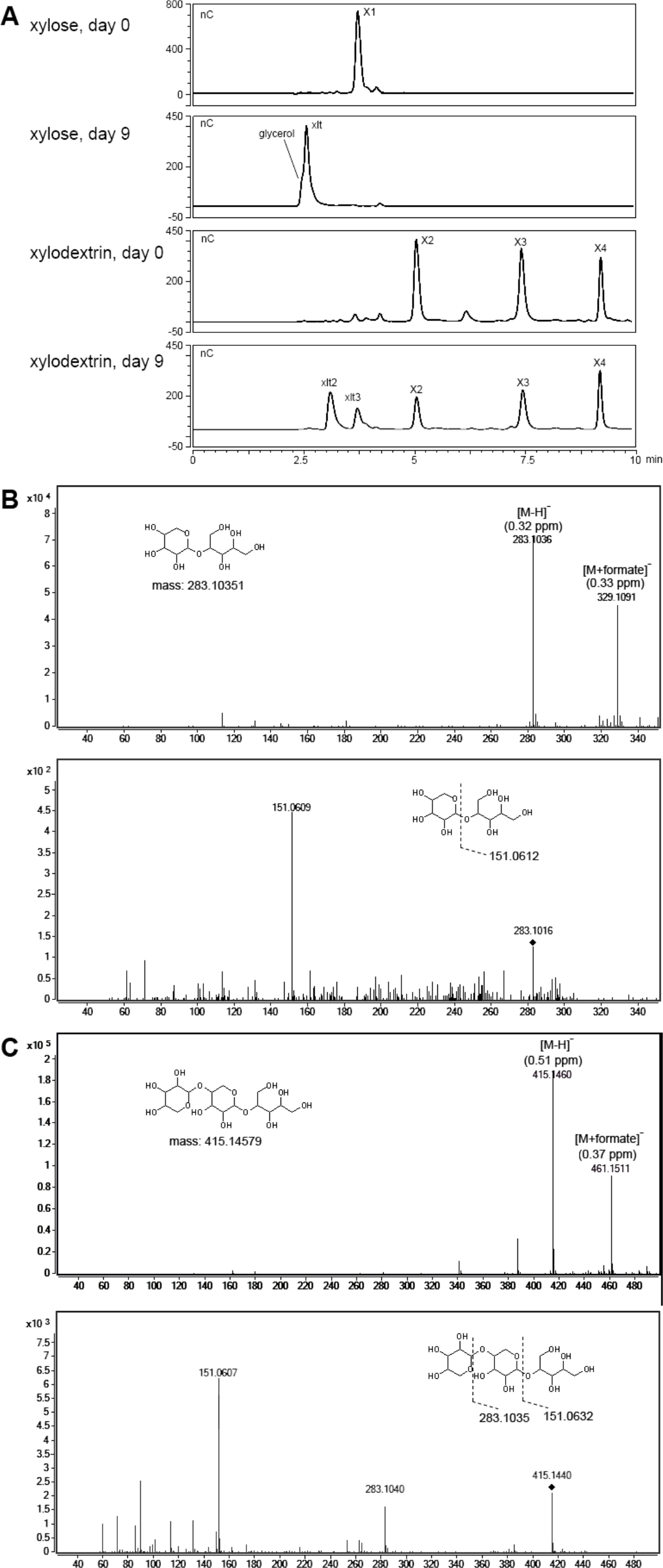
Xylosyl-xylitol oligomers generated in yeast cultures with xylodextrins as the sole carbon source.

**(A)** Carbohydrates from culture supernatants of strain SR8U expressing CDT-2 and GH43-2 (plasmid pXD8.4), resolved by HPAEC. (B) LC-MS and LC-MS/MS spectra for xylosyl-xylitol. High-resolution MS spectra show m/z ratios for the negative ion mode. The deprotonated and formate adduct ions were determined with an accuracy of 0.32 and 0.33 ppm, respectively. The MS/MS spectrum in the lower panel shows the product ion matching the predicted fragment. The parental ion, [xylosyl-xylitol + H]^−^, is denoted with the black diamond mark. (C) LC-MS and LC-MS/MS spectra for xylosyl-xylosyl-xylitol. The deprotonated and formate adduct ions were determined with an accuracy of 0.51 and 0.37 ppm, respectively. The MS/MS spectrum in the lower panel shows the product ions matching the predicted fragments. The parental ion, [xylosyl-xylosyl-xylitol + H]^−^, is denoted with the black diamond mark.

**Figure 2—figure supplement 2.**
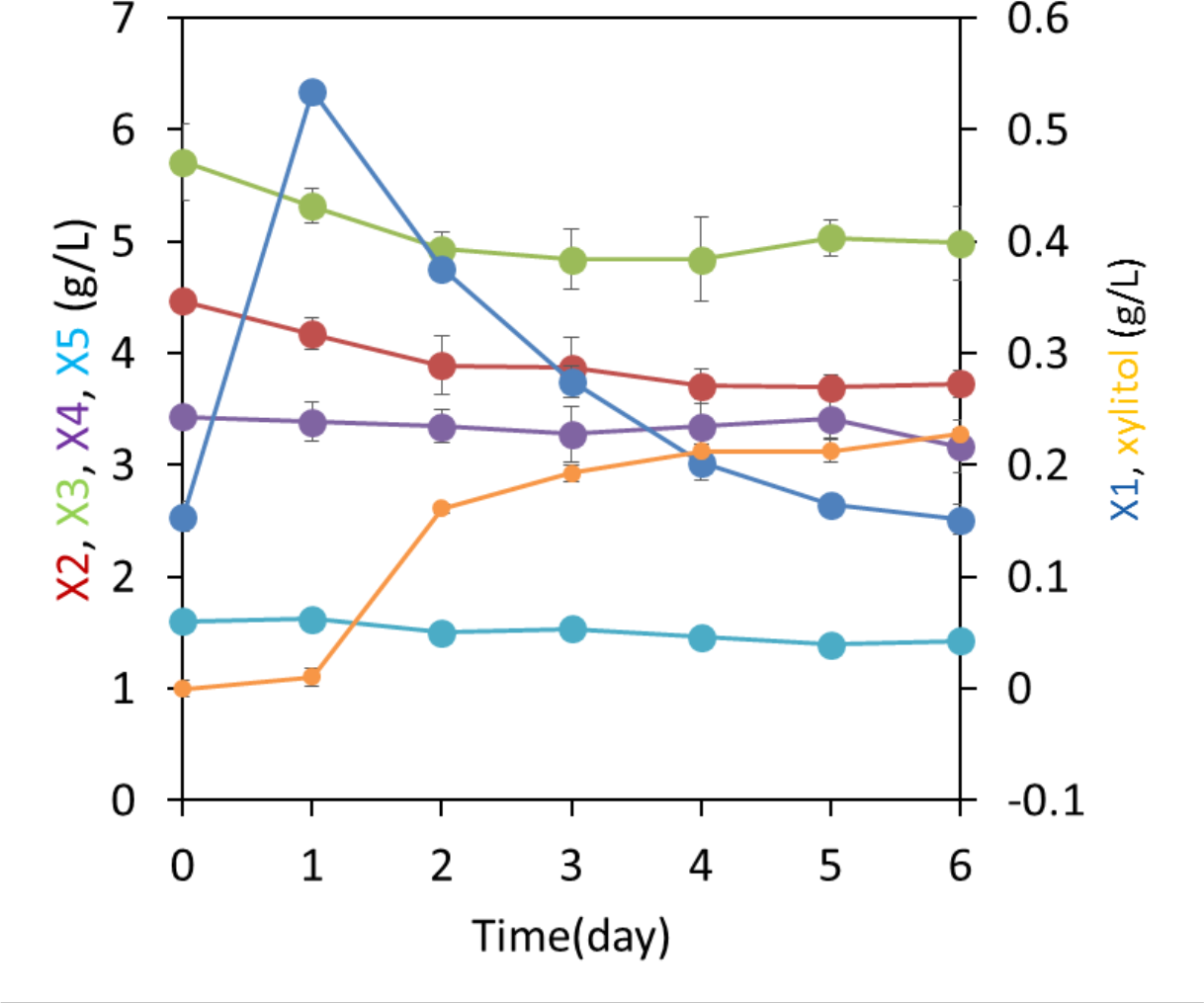
Xylodextrin metabolism by a co-culture of yeast strains to identify enzymatic source of xylosyl-xylitol.

A mixture of a xylose utilizing strain (SR8) with a cell density at OD600 = 1.0 and a xylodextrin hydrolyzing strain (D452-2 expressing CDT-2 and GH43-2 from plasmid pXD8.4) with a cell density at OD600 = 20 were co-cultured in a medium containing 2% xylodextrin. Xylobiose (X2) and xylotriose (X3) decreased, whereas xylose (X1) initially increased. Subsequent X1 consumption correlated with production of xylitol. Notably, xylosyl-xylitol oligomers were not detected, suggesting that the xylodextrin reductase activity was present only in the xylose-fermenting strain expressing XR. Error bars represent standard deviations of biological triplicates.

**Figure 2—figure supplement 3.**
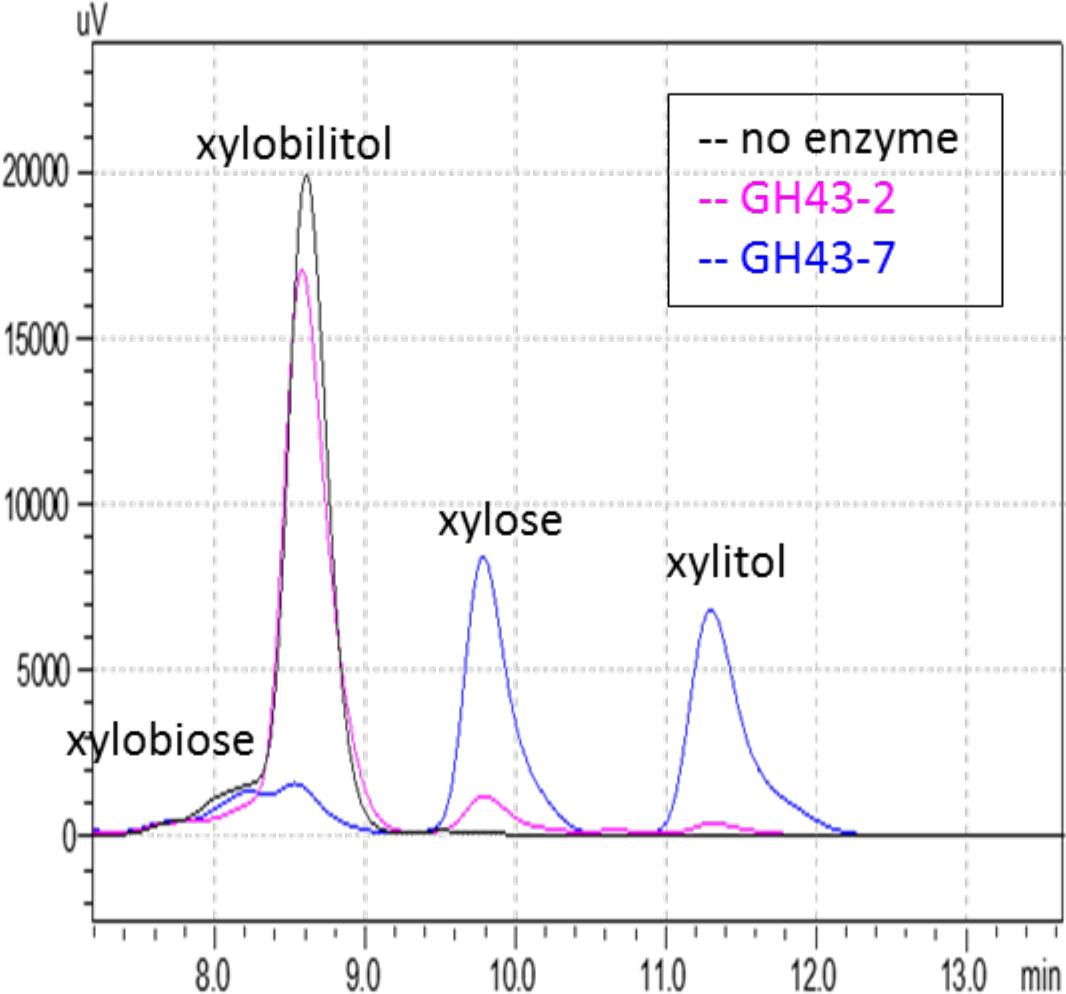
Chromatogram of xylosyl-xylitol hydrolysis products generated by β-xylosidases.

Reaction products from the enzymatic assays in Figure 2D were resolved by ion-exclusion HPLC. Peak areas were used to quantify the concentration of substrates and products at the end of the reaction.

**Figure 3—figure supplement 1.**
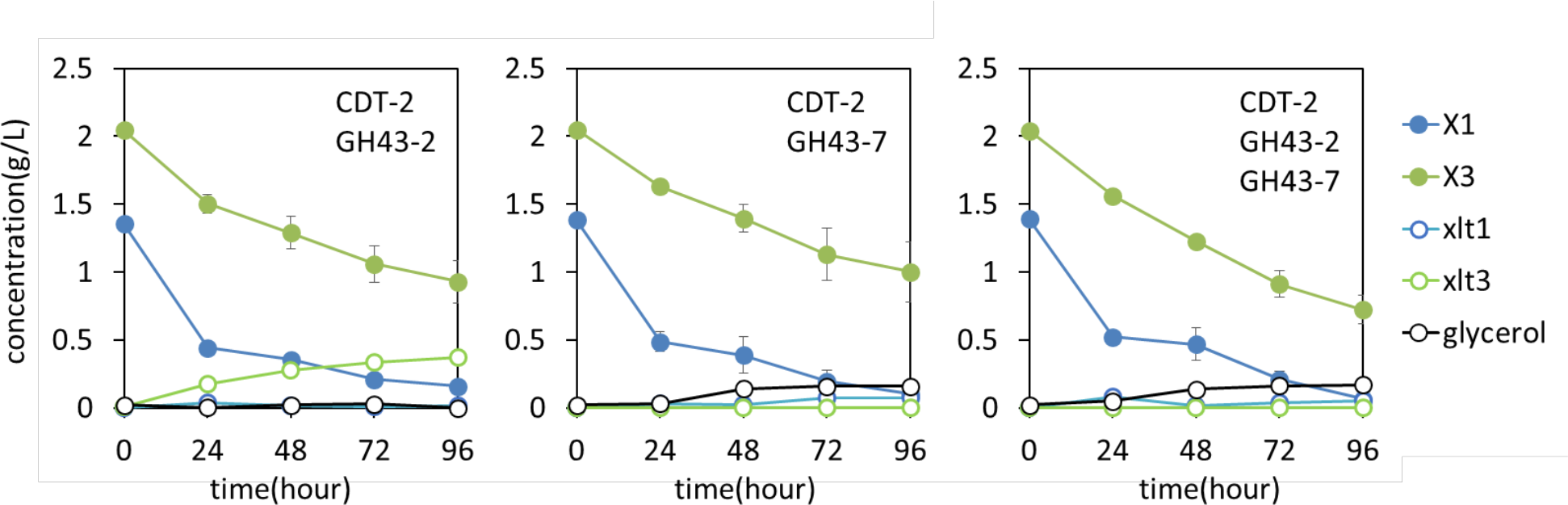
Culture media composition during yeast growth on xylodextrin.

Yeast growth with xylodextrin as the sole carbon source under aerobic conditions with a cell density at OD600 = 20 or under microaerobic conditions with a cell density =80. Yeast strain SR8 transformed with plasmid expressing CDT-2 and GH43-2 (pXD8.4), CDT-2 and GH43-7 (pXD8.6) or all three genes (pXD8.7). All growth experiments were performed in biological triplicate and error bars indicate the standard deviation between experiments.

**Figure 3—figure supplement 2.**
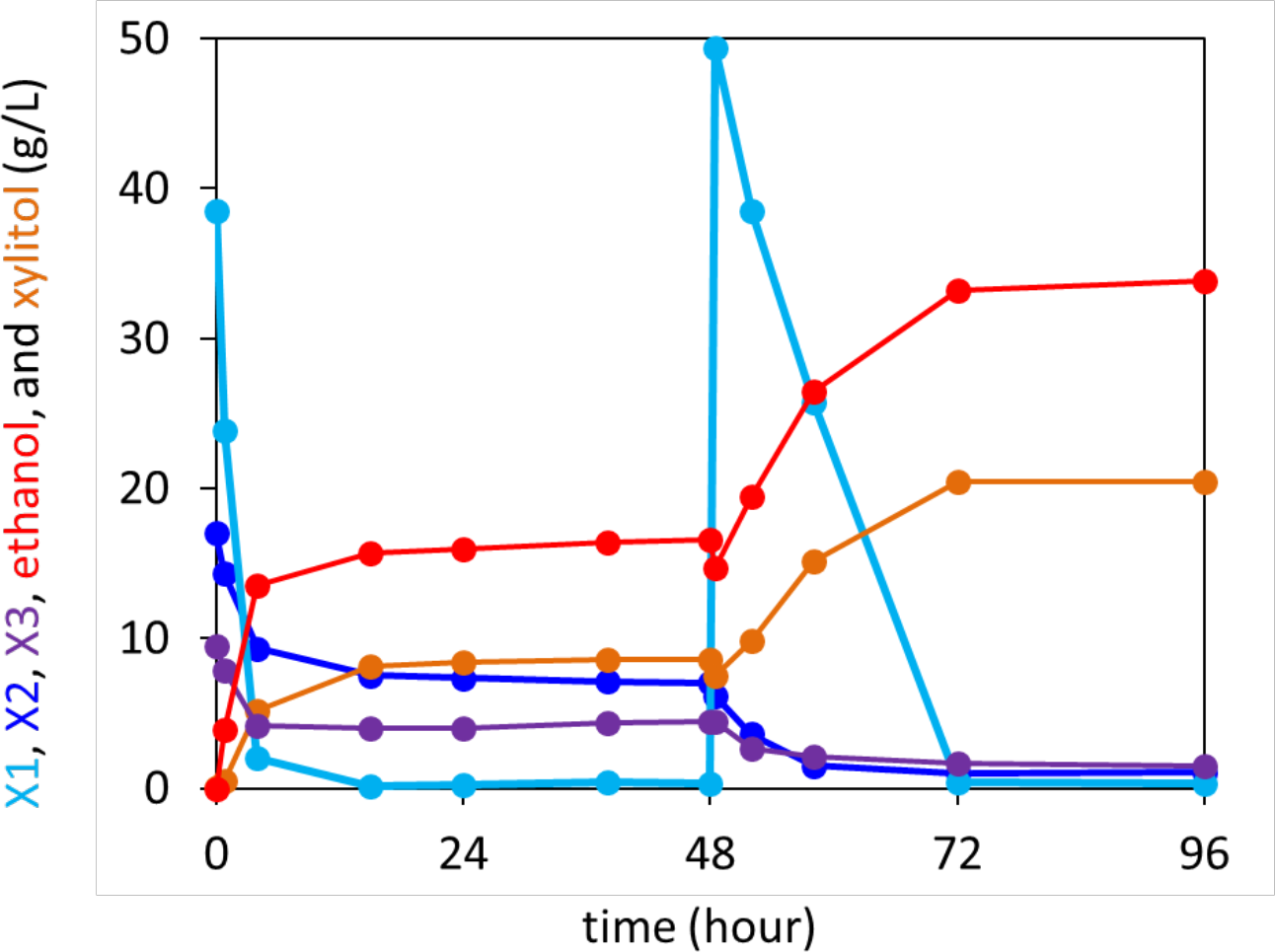
Anaerobic xylodextrin utilization requires the presence of xylose.

Strain carrying the complete xylodextrin pathway grown under anaerobic conditions in oMM containing 4% xylose and 3% xylodextrin. The consumption of xylobiose (X2) and xylotriose (X3) stalled when xylose (X1) was depleted and resumed after supplying additional xylose at hour 48.

**Figure 3—figure supplement 3.**
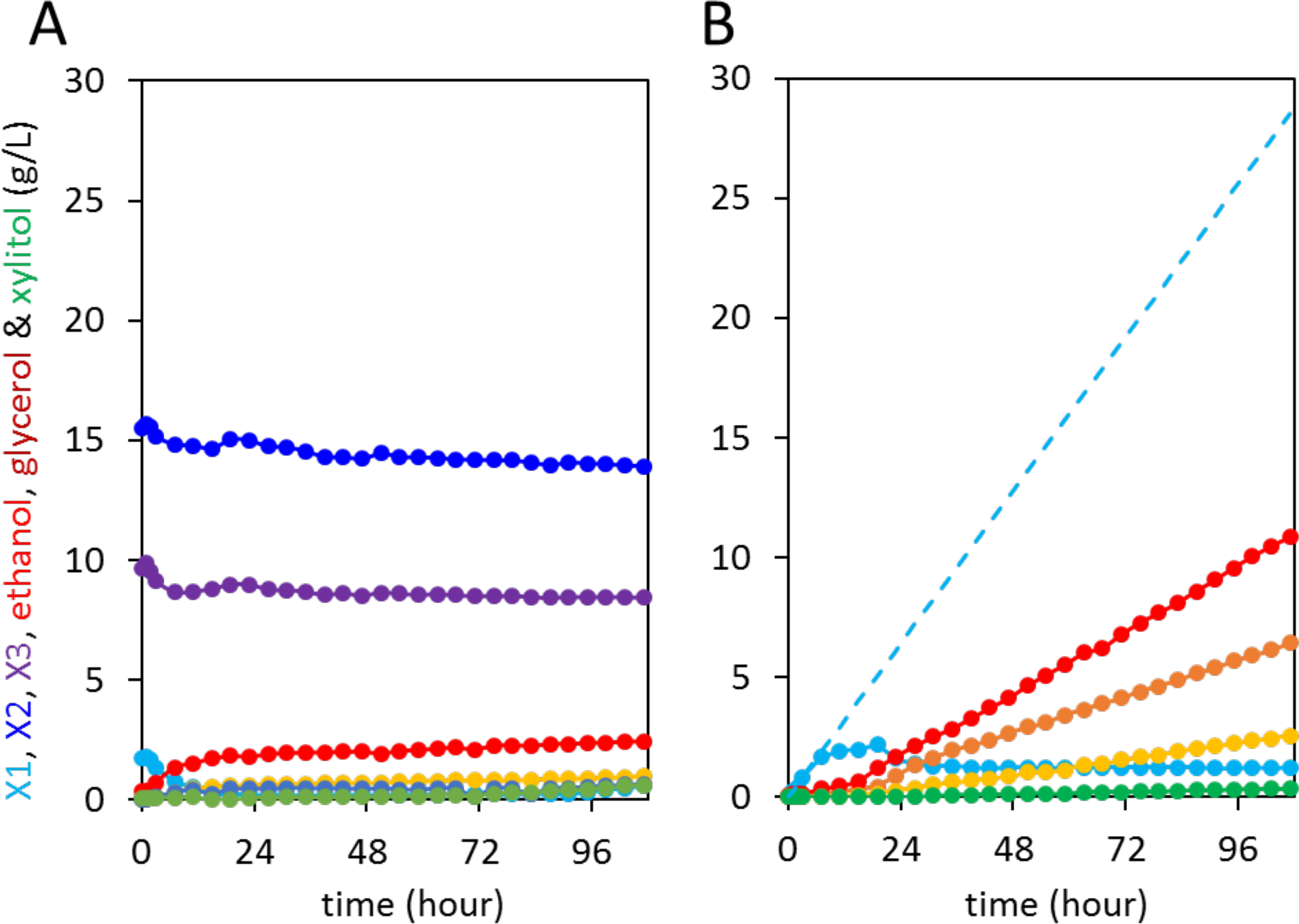
Control anaerobic fermentations with *S. cerevisiae* strain expressing the complete xylodextrin utilization pathway.

Strain SR8 with plasmid pXD8.7 was used at an initial OD600 of 20. Solid lines represent concentrations of compounds in the media. Blue dotted line shows the total amount of xylose added to the culture over time. (A) Fermentation profile of oMM medium containing 4% xylodextrin in the reactor without feeding xylose. (B) oMM medium without xylodextrin in the reactor but with continuous xylose feeding.

**Table S1.** A list of plasmids used in this study.

| Plasmid | genotype and use | use | Ref. |
| --- | --- | --- | --- |
| pRS426_NCU08114 | $P_{PGK1}$-CDT-2 | transport assay | (Galazka, Tian et al. 2010) |
| pRS423_GH43-2 | $P_{TEF1}$-GH43-2 | enzyme purification | this study |
| pRS423_GH43-7 | $P_{TEF1}$-GH43-7 | enzyme purification | this study |
| pRS313_NcXR | $P_{CCW12}$-NcXR | enzyme purification | this study |
| pET302_EcGH43-7 | EcGH43-7 | enzyme purification | this study |
| pET302_BsGH43-7 | BsGH43-7 | enzyme purification | this study |
| pLNL78 | $P_{RNR2}$-SsXK::$P_{TEF1}$-SsXR::$P_{TEF1}$-SsXDH | fermentation | (Latimer, Lee et al. 2014) |
| pXD2 | $P_{RNR2}$-SsXK::$P_{TEF1}$-SsXR::$P_{TEF1}$-SsXDH::$P_{PGK1}$-CDT-2::$P_{TEF1}$-GH43-2 | fermentation | this study |
| pXD8.4 | $P_{CCW12}$-CDT-2::$P_{CCW12}$-GH43-2 | fermentation | this study |
| pXD8.6 | $P_{CCW12}$-CDT-2::$P_{CCW12}$-GH43-7 | fermentation | this study |
| pXD8.7 | $P_{CCW12}$-CDT-2::$P_{CCW12}$-GH43-7::$P_{CCW12}$-GH43-7 | fermentation | this study |

